# Large-scale analyses of short open reading frames yield few if any novel human genes

**DOI:** 10.64898/2026.11.19.767917

**Authors:** Hyun Joo Ji, Steven L. Salzberg

## Abstract

Recent efforts to discover new protein-coding genes in the human genome have yielded a large number of short open reading frames supported by limited experimental evidence supporting translational activity in fragmented parts of those sequences. We examined the 19 highest-confidence predictions made by the TransCODE Consortium, and found that three of them were previously known and thus not novel, while the other 16 could be explained by mechanisms that do not require the assertion of a novel human protein. In several cases, the alternative explanations for the other 16 had already been published. We also analyzed the other 118 predictions and found that most of these, too, could be explained without positing the existence of novel proteins. Our findings show that in the search for new human genes, evidence of transcription and translation can almost always be explained by known mechanisms, and that the predictions of the TransCODE Consortium likely do not include any novel genes.

## Introduction

The scientific community has been working for decades to identify all the genes in the human genome [1], but we have yet to reach a consensus, in part because many aspects of the problem have turned out to be much more challenging than expected [2]. Because each human gene is potentially so important to our understanding of health and disease, great effort continues to go into finding new genes as well as characterizing the functions of known genes more fully. One strategy in recent years has been to examine the vast number of short open reading frames (ORFs) in the genome, in the hopes that previously-undiscovered genes might still be found hidden in these regions. Newer technology allows us to apply high-throughput assays to determine if these short ORFs are transcribed and translated, which are necessary but not sufficient conditions for identifying a sequence as a protein-coding gene.

However, caution must be exercised when searching through large volumes of data to gather evidence for new genes. The human genome has been the object of intensive study for many decades, and efforts to catalog all genes have been under way since the initial publication of the draft human genome in 2001 [3, 4]. During most of the 25 years since, the gene count has declined rather than increased [2], in part because previously-annotated genes turned out to be pseudogenes or other types of false positives generated by computational pipelines. When the first complete human genome was published in 2022, adding nearly 200 megabases of new sequence to the genome [5], 130 new protein-coding genes were reported, all of them paralogs of known genes. This was the only instance in the past two decades in which a large group of clearly authentic new genes was discovered in the human genome. (Some of these additional genes have since been shown to perform essential functions beyond what previously-annotated paralogous genes provided [6].)

Thus when mining a large set of ORFs in the hunt for new protein-coding genes, researchers should be careful to consider more mundane explanations for their data before concluding that they have found a new gene. In particular, although the production of proteins in a cell requires transcription from DNA to RNA and translation into polypeptides, both transcription and translation are “noisy,” which means that evidence of transcription and translation is not sufficient, *per se*, to prove the existence of a gene. To put this in logical terms, if we know that A implies B and we observe B, we cannot infer that A is true.

In this context, we here evaluate the findings of a recent study by Deutsch *et al*. [7], who conducted an extensive analysis of 7,264 previously-reported [8] non-canonical open reading frames (ncORFs) in the human genome, and determined based primarily on peptide detection from >95,000 proteomics experiments that 137 of these ncORFs should be characterized as either novel proteins or as “peptideins,” which they described as “microproteins that have indeterminate potential as functional proteins.” They divided these into four groups, labeled as Tiers 1A, 1B, 2A, and 2B, and argued that the 18 ncORFs in Tier 1A, the highest-confidence group, were particularly likely to represent functional proteins. They also noted that three of those 18 Tier 1A sequences were recently added to the GENCODE [9] human annotation database as protein-coding genes.

As an aside, we note that the term “peptideins” has a nearly identical meaning to the term “translon,” which was proposed as a general term in 2025 [10] by a large consortium of scientists (including some of the authors of [7]). A translon is any segment of DNA that is translated by the ribosome. There is no need for yet another specialized term, so in the text below we use the term translon interchangeably with peptidein.

## Results

We have carefully inspected all 18 of the Tier 1A ncORFs, plus one from Tier 1B that Deutsch *et al*. specifically highlighted as a novel protein, and found the evidence lacking for nearly all of them. In our detailed analysis of these 19 ncORFs below, we point out several of them had already been analyzed and published by our own lab [11], where rather than claiming them to be novel protein-coding genes, we explained that they are much more likely to represent novel isoforms of existing protein-coding genes. We also applied the pipeline we previously described [11] to all of the Tier 1A, 1B, 2A, and 2B upstream and downstream ORFs (excluding c6norep53, which was reported in [7] to be a pseudogene fragment). Our results indicate that many more of the 137 ncORFs can be explained as protein-coding exons of existing genes, and at least one of them was previously annotated as such in RefSeq [12].

**Table 1** lists the 19 highest-confidence ncORFs from [7], and **Supplementary Table S1** lists the amino acid translations of those ORFs. All were classified as Tier 1A except for c2riboseqorf47 from Tier 1B. Our analysis shows that these ncORFs fall into four groups, which we describe in detail below. The first group (3 candidates) is a set of sequences that were erroneously listed in [7] as ncORFs but that are actually already known or already predicted to be protein-coding genes, so we begin our analysis with that group. The second group (6 candidates) includes ncORFs that were previously published as coding exons from alternative splice variants of the gene containing them. These alternative splice variants provide a simple explanation for any peptide evidence, which is much more plausible than positing the existence of entirely new, very short human proteins. The third group (6 candidates) comprises ncORFs that appear to represent translational noise; i.e., an upstream start codon triggers translation that does not produce a functional protein. The fourth and last group (4 candidates) includes ncORFs that are pseudogenes and known functional RNAs, or otherwise have no evidence that they encode functional proteins.

**Table 1.**
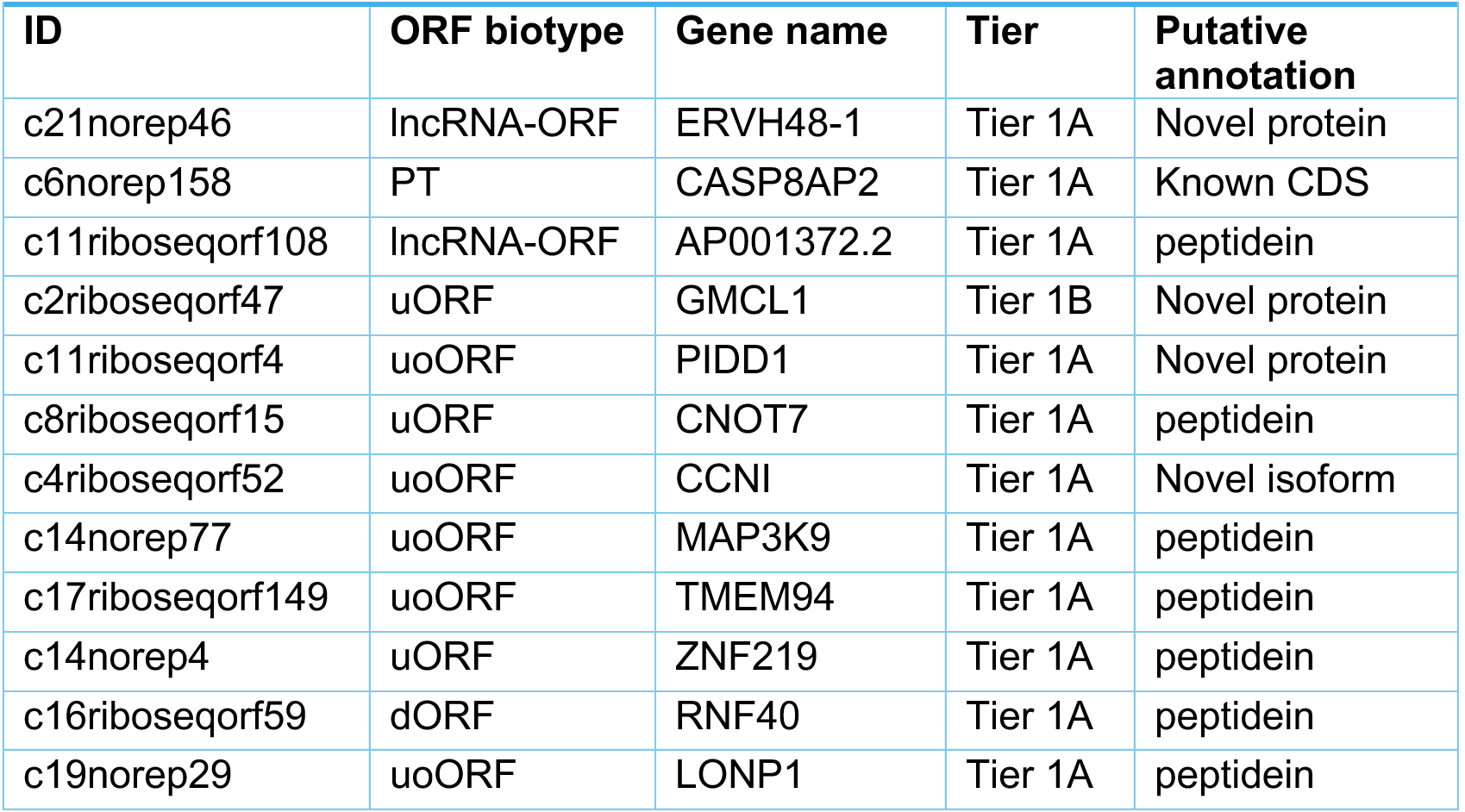
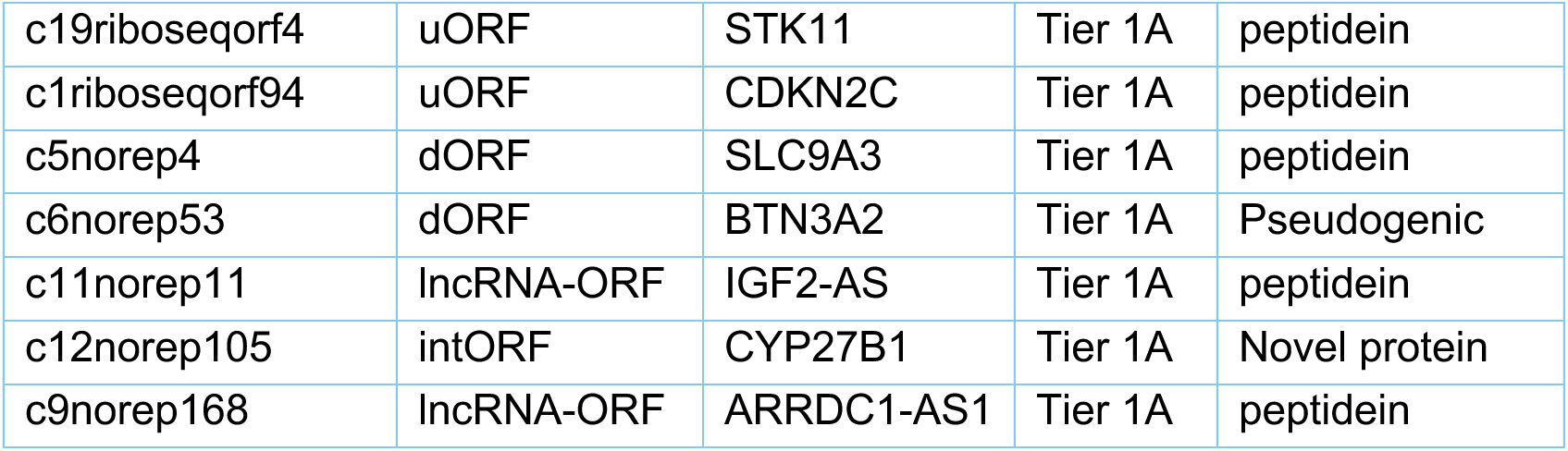
Highest-confidence peptideins and novel proteins from Supplementary Table 12 in Deutsch et al. **[7]**. This table contains all 18 ORFs from Tier 1A plus one ORF from Tier 1B. Column 2 classifies the ORFs based on where they are positioned within the coding sequence of the gene shown in column 3: uORF, upstream ORF; intORF, ORF that is internal to the coding sequence; lncRNA-ORF, ORF within a lncRNA gene; PT, known protein-coding gene; uoORF, upstream ORF that overlaps the coding sequence; dORF, downstream ORF. Column 5 shows the annotation proposed in **[7]**, where “peptidein” was defined as a “confirmed protein molecule of indeterminate consequence.”

### Group 1: known protein-coding genes reported as peptideins

**C21norep46**, a non-canonical ORF in ERVH48-1, was reported by [7] as one of three ncORFs (in addition to c12norep105 and c11riboseqorf4) in their list of peptideins that have recently been upgraded to protein-coding genes in the GENCODE database [9]. Although they list c21norep46 as a lncRNA-ORF (*i.e*., a protein-coding region found within a long noncoding RNA), this sequence has long been known as ERVH48-1, which encodes Suppressyn, a placenta-specific,160aa protein derived from a retroviral envelope protein. ERVH48-1 was first reported as a protein-coding gene, in 2013 [13], along with evidence for its function, so it is not a novel finding in [7], and it was also incorrect to report it as a noncoding ORF in a previous paper by some of the same authors [8]. ERVH48-1 is also annotated as a protein-coding gene in MANE and CHESS [14], including CHESS release 2 published in 2018 [15]. So the report that c21norep46 is a lncRNA was simply incorrect.

**C6norep158** was reported in [7] as a Tier 1A peptidein, but they also describe it as a “known CDS” in the gene CASP8AP2. This gene has been annotated in both RefSeq [12] and CHESS for at least a decade, and it is the only one of the 137 ncORFs in Tiers 1A, 2A, 1B, and 2B that is described in [7] as a known protein. It is unclear why CASP8AP2 was included in the original list of 7,264 noncoding ORFs collected by a subset of the same authors [8], but it is a well-known protein-coding gene, and its inclusion in a set of ncORFs appears to be an error.

**C11riboseqorf108** occurs within a lncRNA annotated in GENCODE as LIPT2-AS1, reported as a peptidein of length 349aa (Supplementary Table S1), making it the longest of the putative novel proteins reported in [7]. However, this annotation is not new: it was first annotated as a protein-coding gene in one of the 2001 papers reporting the initial sequence of the human genome [4], when it was assigned GenBank accession EAW74941.1.

This sequence is not annotated as a protein in the current versions of RefSeq [12], GENCODE [9], and CHESS [14], but it was relatively recently included in both RefSeq and CHESS as a protein, with identifiers NP_001306169.1 and CHS.9489 respectively. The predicted protein was removed from RefSeq in 2020 and changed to a lncRNA, which agreed with GENCODE. In 2023, it was changed to a lncRNA in CHESS as well, based on the agreement between RefSeq and GENCODE that the sequence was a lncRNA. The previously annotated protein from CHESS (release 2.2) is 392aa in length and contains 100% of c11riboseqorf108; the only difference is that CHS.9489 uses an upstream start codon that makes it longer by 43aa. The protein has highly significant matches to annotated proteins in many primates, mammals, and more distant species; the evidence is clear that this is a protein-coding gene, not merely a peptidein.

### Group 2: peptideins that can be explained as coding exons from alternative splice isoforms

Deutsch *et al*. highlighted **c2riboseqorf47** as a particularly strong case for a novel protein, writing: “our development of the ORBL method to analyse ORF evolutionary constraint, our assessment of CRISPR-based functional inference and our prioritization of HLA immunopeptide support were central to the establishment of c2riboseqorf47 as a protein-coding gene” [7].

Despite this claim, a much simpler explanation, previously published in 2024 [11], is that c2riboseqorf47 contains an unannotated coding exon of GMCL1, created by splitting the first exon into two exons separated by a short intron. **Figure 1** illustrates the position of this putative coding exon, which contains 9aa (METVASAAT) and is then linked to the next exon through an intron whose splice sites are supported by spliced RNA data, as reported previously [11]. The putative 19aa protein encoded by c2riboseqorf47 is defined by extending the coding sequence of that first exon until it reaches the first downstream stop codon, shown in the lower panel of **Figure 1**.

**Figure 1.**
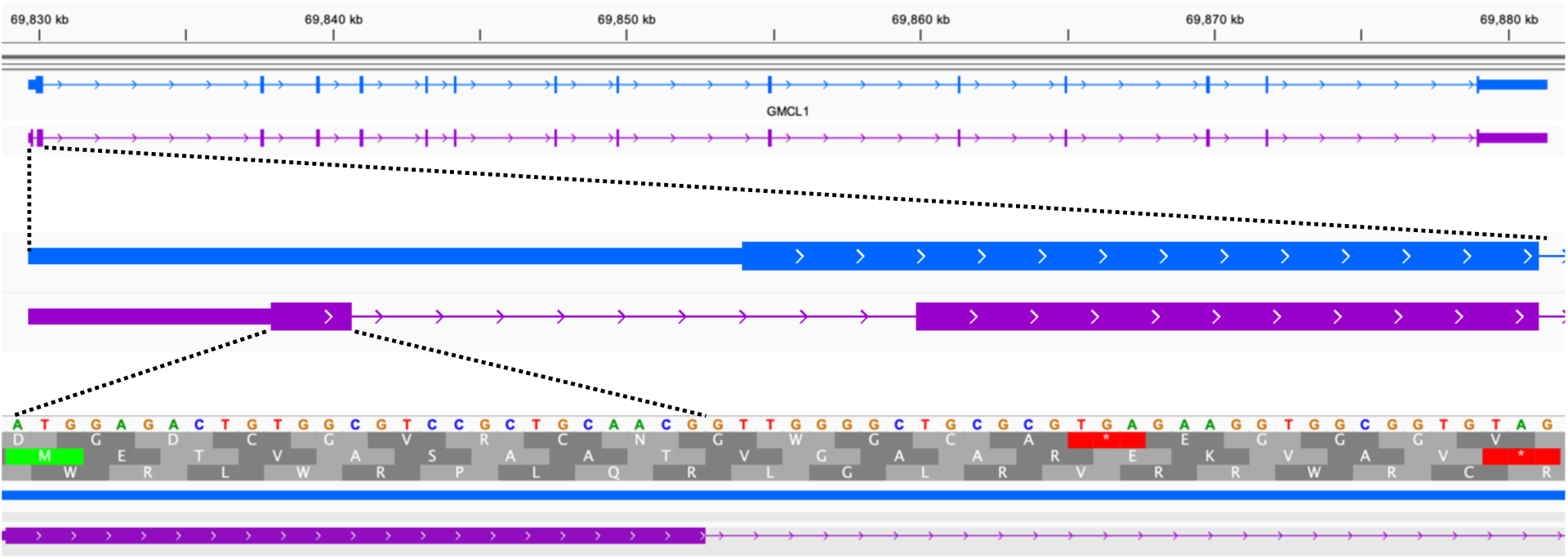
Structure of the GMCL1 gene with an alternative first exon. The top panel shows the canonical sequence in blue and the alternative transcript in mauve, with the canonical exon 1 split into two exons, separated by a short intron. The middle panel shows a zoomed-in view of the beginning of the transcript with its new intron. At bottom, the view is expanded further to show the 9 amino acids of the first exon (METVASAAT) and the donor splice site (GT) following it.

Thus rather than claiming an unusually short, entirely novel gene of just 19aa that resides in the 5’ UTR of another gene, we can explain the evidence for c2riboseqorf47 by the presence of a coding exon in a GMCL1 splice variant. **Figure 1** shows how this transcript replaces the first 19aa of the canonical (MANE) isoform with 9aa from c2riboseqorf47, but otherwise maintains the reading frame of GMCL1 and encodes a protein that shares the remaining 496 amino acids. Worth noting here is that the ribosome profiling data cited in support of c2riboseqorf47 is equally well explained by the translation of this novel exon, which begins at the same start codon. We confirmed that this coding exon is not currently annotated in GENCODE, RefSeq, or CHESS, consistent with our previous characterization of it [11] as a potentially novel protein coding exon.

**C11riboseqorf4** is a 171aa ORF that is upstream of and overlapping with the coding sequence of PIDD1. Deutsch *et al*. claim that this is an entirely new human protein-coding gene, in part because peptides from this uoORF (upstream overlapping ORF) have been identified in mass spectrometry experiments from human tissues and cell lines. Their proposed new gene uses a start codon (ATG) that is 31bp upstream of the canonical start codon for PIDD1, thereby using a different open reading frame and creating a protein sequence that shares no similarity with the 910aa protein produced by the canonical sequence of PIDD1. Deutsch *et al*.’s predicted short protein does not show similarity to any other known proteins.

We analyzed this same uoORF in our previous study [11], in which we found RNA-seq evidence suggesting that the uoORF might instead be a protein-coding exon belonging to an unannotated transcript of PIDD1. Here we re-analyze the splicing evidence in this region, noting first that c11riboseqorf4 consists of two partial exons that overlap exons 2 and 3 of the MANE isoform (**Figure 2**, with the uoORF shown in black near the top). We identified three distinct splice variants (labelled 1–3 in **Figure 2**), each of which begins at the ncORF’s start codon and uses splice junctions that are supported by GTEx RNA-seq data. In each case, the variant preserves ≥ 80% of the annotated coding sequencing of PIDD1. For example, c11riboseqorf4_c1 uses the exon 2 splice donor and skips exon 3, splicing into exon 4 at an acceptor site that restores the canonical reading frame. The other two candidates use different GTEx-supported donor sites upstream of the canonical donor site. Both produce isoforms that restore the canonical PIDD1 reading frame without skipping exon 3.

**Figure 2.**
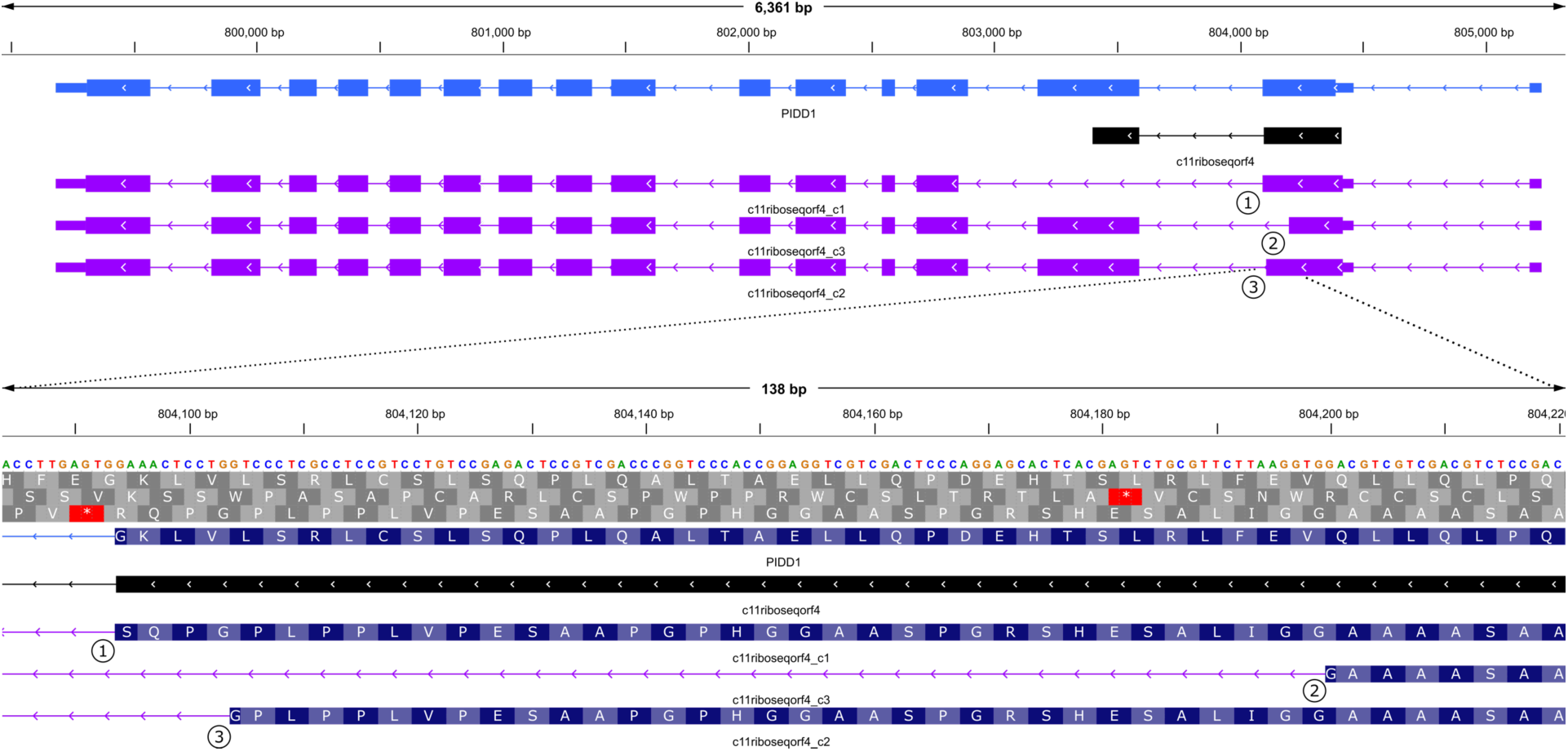
Structure of the PIDD1 gene with three putative splicing variants supported by RNA-seq evidence. The top panel shows the canonical (MANE) sequence in blue, the ncORF from Deutsch *et al*. in black, and the alternative transcripts described here in mauve. The first coding exon of each splice uses the same start codon as c11riboseqorf4, the ncORF in question, and each is followed by an intron that has strong RNA-seq support. The bottom panel provides a close-up of these exons, illustrating that they share peptide sequences with c11riboseqorf4, which in turn would produce similar mass spectrometry evidence.

As with c2riboseqorf47, the presence of an upstream ATG on the transcripts of PIDD1 allows the ribosome to occasionally begin translation at an earlier start codon, producing peptides that may very well appear in mass spectrometry and Ribo-Seq experiments. The presence of these peptides is not sufficient evidence to claim that they are part of a novel short protein, especially when, as here, the peptide evidence reported in [7] could be explained more mundanely by any of the alternative isoforms shown in **Figure 2**.

**C8riboseqorf15** is an upstream ORF starting in the first exon of CNOT7, which has a noncoding first exon in its canonical transcript. The peptidein reported in Deutsch *et al*. has a short (39aa) and highly repetitive sequence (Supplementary **Table S1**). This ORF too was previously reported in [11], where we described it as a novel protein coding exon from an alternative splice variant of CNOT7, a 285aa protein.

For the putative peptidein c8riboseqorf15, the first 36aa are translated from the first exon, and its final three amino acids (VHK) come from the second exon, ending at a stop codon. In our study, we reported that the upstream ORF sequence starts at the identical site and includes the same initial 36aa sequence. Specifically, we described a putative novel isoform that then skips exon 2 and continues with exon 3, where the protein translation remains in-frame, encoding a protein nearly identical to the canonical version of CNOT7, except for the N-terminus, where the initial 36aa are replaced by the first 36aa from the sequence shown in Supplementary **Table S1**. We used AlphaFold2 to predict the structure of the resulting protein, and found that it received a pLDDT score of 86.3, quite high but slightly lower than the MANE isoform, which had a score of 90.7.

The Ribo-seq evidence used to elevate c8riboseqorf15 to Tier 1A provides equally good support for the translation of the alternative CNOT7 isoform that we published earlier [11]. Thus rather than positing a new and unusually short human protein-coding gene, an alternative protein-coding transcript provides a simpler and more plausible explanation for evidence of translation from this upstream ORF.

**C4riboseqorf52** is a uoORF in the gene CCNI. As with the ncORFs mentioned above, this uoORF was also previously reported as a putatively novel protein-coding exon, rather than an entirely novel gene [11]. The sequence of c4riboseqorf52 comprises 59aa (Supplementary **Table S1**), which begins at an alternative upstream start codon and ends in the middle of exon 2. Our alternative isoform, in contrast, uses the same first 24aa (encoded in exon 1), but the first exon is spliced into downstream exon 3, matching the canonical CCNI protein sequence from that point on (Supplementary **Figure S1**). Thus our previously-reported isoform encodes a protein that replaces the first 38aa of the canonical sequence of CCNI with the first 24aa of c4riboseqorf52, and otherwise matches CCNI. Worth noting is that a full-length cDNA published in 2004 [16] used the same upstream start but skipped exon 2 and matched the remaining canonical sequence of CCNI, identically to what we described in [11]. Our re-analysis here also suggested several additional alternative isoforms at this locus, some of which skip only part of exon 2, while others skip both exons 2 and 3. All of these isoforms use the same start codon as c4riboseqorf52, and any of them would yield similar peptide evidence in Ribo-Seq or mass spectrometry data.

**C14norep77** is a uoORF of length 161aa (Supplementary **Table S1**) in the gene MAP3K9, whose canonical isoform is far longer, 1104aa. This upstream, overlapping ORF starts just 16bp upstream of the annotated start codon for MAP3K9, and produces a short translated sequence that ends 3aa into the second exon. The MAP3K9 gene has 12 exons in human, and the alternative translation of exon 1 (corresponding to c14norep77) has been annotated in chimpanzee (GenBank accession XP_016781765.3) as a splice variant of a full-length isoform (1107aa) of the chimp protein.

In our previous analysis [11], we excluded c14norep77 from the ncORFs we examined because it was not reproducibly detected across the Ribo-seq studies considered in [7]. However, because this ncORF is listed as a peptidein in [7], we analyzed it further here. This uoORF consists of parts of two exons, overlapping exons 1 and 2 of the MANE isoform (**Figure 3**, uoORF shown in black near the top). Using splice sites found in GTEx RNA-seq data, we identified two alternative transcripts that contain ORFs which use the same start codon as c14norep77, but generate translations that match the canonical MAP3K9 reading frame (**Figure 3**). One of the alternative isoforms (labeled 1 in **Figure 3**) contains a coding region that replaces just 30 aa of the canonical protein’s N-terminus with 13 aa from c14norep77, and the resulting protein sequence is 98.5% identical to the MANE isoform. Thus the translation evidence reported in [7] for c14norep77 can be explained more simply as the product of alternative isoforms of MAP3K9.

**Figure 3.**
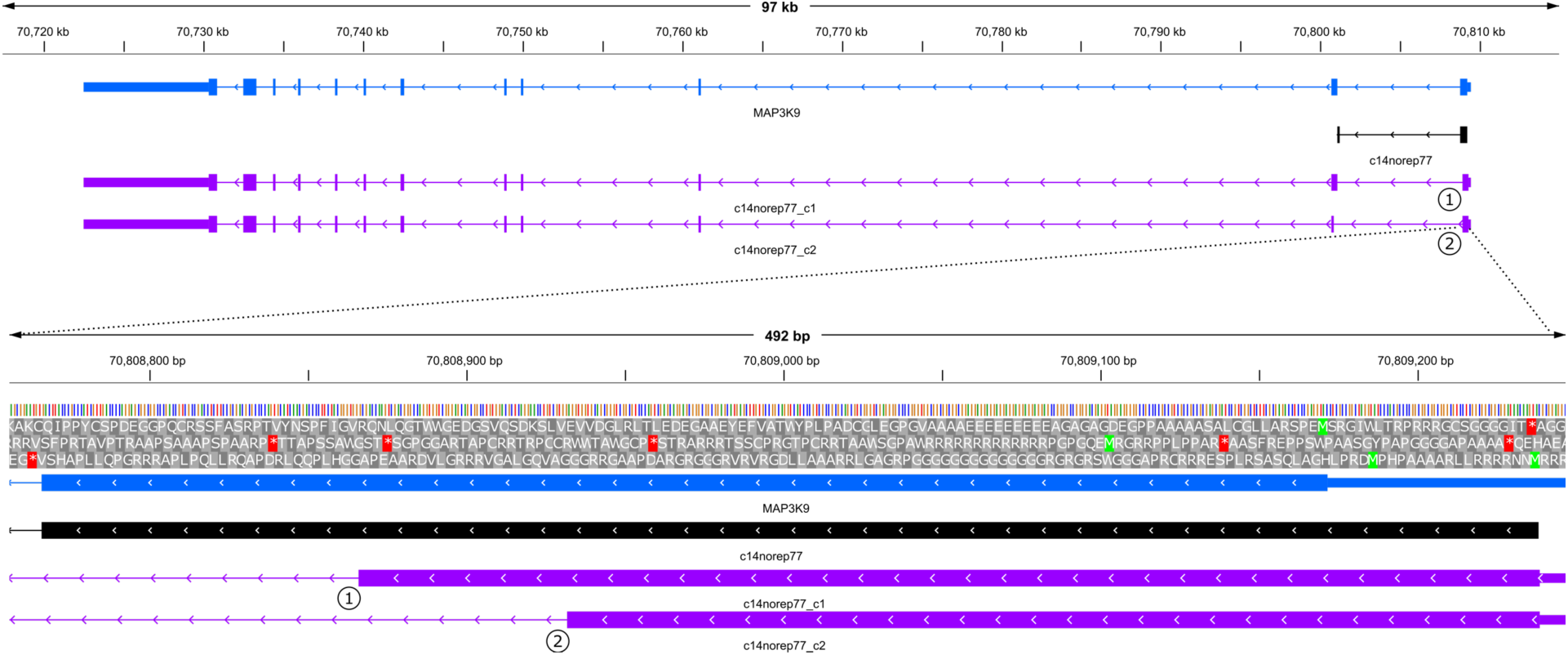
Structure of the MAP3K9 gene with two putatively novel exon candidates supported by RNA-seq evidence. The top panel shows the canonical (MANE) sequence in blue, the ncORF in black, and alternatively spliced transcripts in mauve. Two alternative transcripts are shown, whose first exons illustrate the sequences derived from c14norep77. The bottom shows a close-up of exon 1, and illustrating how the alternative isoforms using different splice sites from the ncORF. Both transcripts use an earlier donor site than the ncORF; alternative transcript 1 uses the same splice acceptor as the MANE transcript, whereas alternative transcript 2 uses an different acceptor site within the second coding exon.

**C17riboseqorf149** is an upstream, overlapping ORF in TMEM94, encoding a translon of 84aa (Supplementary Table S1). The canonical sequence of TMEM94 is 1356aa. The uoORF begins at a start site slightly upstream of the canonical TMEM94 start and then spans three exons, which correspond to exons 2–4 of the MANE transcript (Figure 4). We found clear RNA-seq evidence supporting an alternative transcript that utilizes an acceptor site in exon 3 located 16bp downstream of the canonical one (see transcript variant (1) in Figure 4). This junction is supported by both short- and long-read RNA-seq data, and the resulting translation is nearly identical to the canonical TMEM94 protein; the only difference is that it replaces the first 13aa of the canonical protein with 31aa from the ncORF. We also identified multiple other splice variants that use the same start codon as c17riboseqorf149 but then use alternative splice sites that restore the reading frame to match the canonical one (variants 2–7 in Figure 4). Thus as with the previous cases, the peptide evidence for this ncORF can be explained by any of several alternative splice variants of TMEM94.

**Figure 4.**
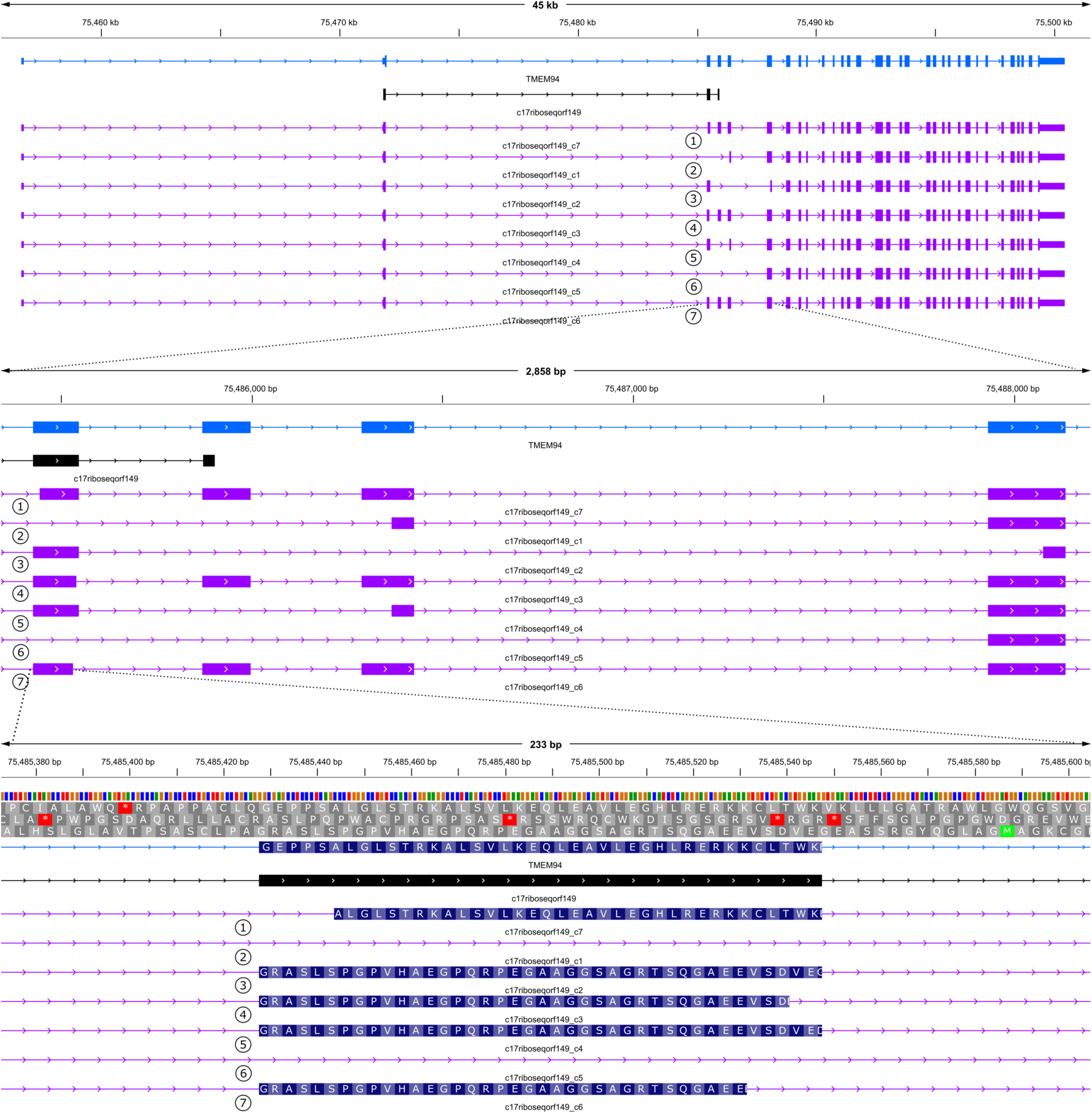
Structure of the TMEM94 gene, showing seven distinct alternative splicing patterns and five putative exon candidates. The top panel shows the canonical (MANE) transcript in blue, the ncORF in black, and the alternative transcripts in mauve. The middle panel shows the splicing patterns near exons 3–7. Four alternative splice variants (labelled 2, 3, 5, and 6) skip the fourth exon entirely, where the ncORF ends at a stop codon, and then splice into a later exon that restores the reading frame. The bottom panel shows a zoomed-in view that illustrated how transcript 1 uses a different acceptor site for exon 3 that matches the canonical reading frame. The remaining alternative splice variants (4 and 7) restore the canonical frame using alternative donor sites on the 3’ end of exon 3.

### Group 3: Likely translational noise from non-functional ncORFs

**C14norep4** is a ncORF of length 80aa in the gene ZNF219, which is 722aa long. This uORF begins in the first exon of ZNF219, which is entirely noncoding, and ends at a stop codon just 8aa into the second exon. In rhesus macaque, ZNF219 is longer, 967aa, and the first 72aa of c14norep4 are 97% identical to positions 80-153 of the macaque protein (GenBank accession XP_077796629). Thus this ncORF is annotated as an alternative upstream coding exon in the ZNF219 gene in macaque. However, translations beginning in the first exon do not form a consistent open reading frame with the remaining exons in the human version of the gene. The peptide evidence reported in [7] can be explained as translation activity that begins in the first exon but does not produce a functional protein. Thus c14norep4 can be categorized as a translon; *i.e.*, a translated fragment of the genome that is not part of a gene [10].

**C16riboseqorf59** appears as a downstream ORF in RNF40. Its translation is 194aa long (Supplementary Table S1), while the canonical sequence of RNF40 is 1001aa. This translon appears in a UTR that is >2200bp in length, and its hypothesized start coding is >1000bp downstream from the stop codon of RNF40. The translated sequence has no homology to any known protein and no known function. It is possible that the peptide evidence identified in Deutsch *et al*. is a genuine match that represents translational activity [17]; *i.e*., that in some cases the ribosome engages in the 3’ UTR of RNF40, but there is no other evidence that this downstream ORF encodes a separate protein.

**C19norep29, c19riboseqorf4, and c1riboseqorf94** are all upstream ORFs in known protein-coding genes (LONP1, STK11, and CDKN2C respectively). In all three, translation produces a protein sequence that is much shorter than the known protein and that has no homology to proteins in any other species, making it highly unlikely that these ORFs encode functional proteins. In all three cases, the presence of a start codon upstream of the canonical start could easily cause the ribosome to bind and translate in a different reading frame [18], producing peptides that may be detected in mass spectrometry experiments [17]. Such evidence falls well short of what is needed to prove the existence of a novel, functional human protein.

**C5norep4** is a 113aa downstream ORF (dORF) in SLC9A3. It has no homology to any known protein and may be simply a long stretch of DNA without a stop codon. As with other cases described above, it is possible that the ribosome may occasionally translate this ORF, yielding nonfunctional peptides that can be detected in proteomics assays.

### Group 4: ncORFs that are pseudogenes and lncRNAs

**c6norep53** is a downstream ORF in BTN3A2 that is described by Deutsch *et al*. as a pseudogenic insertion. In other words, this is not a peptidein or a hypothesized novel protein, but merely the remnant of an ORF from a pseudogene. It is not clear why they included it in their list of ncORFs, because their own study called it a pseudogene. As an aside, we note here that BTN3A2 includes two noncoding 3’ exons, with an unusually long last exon, and if that annotation were adjusted, the pseudogene would fall outside the transcript.

**c11norep11,** reported as a peptidein of length 131aa, is in IGF2-AS, a lncRNA transcript that falls within the gene IGF2 on the opposite strand. The open reading frame for c11norep11 starts in the first exon of IGF2-AS and continues into the second exon of this 3-exon lncRNA. However, IGF2-AS was reported as a protein-coding gene (and given the name IGF2-AS) in a large-scale analysis of human transcripts published 2002 [19], and again in a 2006 paper describing the sequence and annotation of human chromosome 11 [20]. Subsequent research showed that IGF2-AS is an antisense RNA gene that has been extensively studied in dozens of publications (e.g., [21]). It is believed to function through its RNA-level regulation, and translation is reported to trigger nonsense-mediated decay rather than producing a functional protein. No homologous protein sequence is found in any other species, further undermining support for the claim that this is a protein-coding gene. Thus although the translation of c11norep11 was annotated as a protein-coding gene decades ago, more recent work shows that its annotation as a lncRNA (in CHESS, GENCODE, and RefSeq) is likely to be correct.

**c12norep105** is an open reading frame in the middle of CYP27B1 that occurs out of frame from the standard protein sequence for that gene. This internal ORF spans 594 bp (198aa) that begins in exon 3 and ends in exon 6. While it is somewhat unusual for an ORF to span this length by chance, a BLAST search of the translated amino acid sequence returns no matches to any other species, so there is no evidence that this 198aa represents an evolutionarily conserved protein. As reported in [7], this ncORF has been promoted to a protein-coding gene by the GENCODE database, but it is not contained in RefSeq or CHESS, and [7] does not provide any details specific to this ncORF. We note that the canonical CYP27B1 gene encodes a 508aa protein that has been extensively studied in hundreds of publications, but no previous reports have claimed that this internal ORF represented a separate, functional protein.

**C9norep168** is a 179aa ORF inside a lncRNA, ARRDC1-AS1. This lncRNA is a 3-exon noncoding sequence annotated in between (and on the opposite strand of) two protein coding genes, ARRDC1 and EHMT1. (Its name is taken from one of those genes.) To create this ORF, an intron missing from all three current annotation databases was added to the lncRNA, between amino acids 26 and 27 in the hypothesized protein sequence. The resulting protein sequence has no homology to any protein outside primates, and those primate annotations are all described as uncharacterized proteins. This does not appear to be a functional protein.

### Additional analyses

In addition to the 19 high-confidence peptideins discussed above, we examined **c14norep128**, a downstream ORF in TNFAIP2 that was described as a “novel” CDS isoform and placed in Tier 1B by Deutsch *et al*. [7]. C14norep128 encodes a 93aa translon (Supplementary **Table S1**), while the canonical TNFAIP2 protein is 654aa. We identified an alternatively spliced transcript that is already annotated in RefSeq, CHESS, and GENCODE and that explains the peptide evidence for this dORF. The alternative transcript, which is supported by extensive RNA-seq data from GTEx including both short and long reads, uses an intron that originates in the 12^th^ and last exon, 6bp upstream of the canonical stop codon, and extends into the ncORF (Supplementary **Figure S2**). This splice variant skips the first 29aa of the ncORF and retains the remaining 64aa. Given that 69% of c14norep128’s coding sequence is already annotated as a coding exon, it is questionable whether it should have been reported as novel, and in any case the peptide evidence for c14norep128 can be explained more simply by these known isoforms.

We also note that fifteen of the 137 ncORFs classified as peptideins in [7], including c2riboseqorf47, c8riboseqorf15, and c4riboseqorf52 analyzed above, were previously characterized as protein-coding exons in our earlier work [11]. In that study, we used RNA-seq evidence and AlphaFold2-predicted structure scores to identify alternatively spliced transcripts of known protein-coding genes that could provide simpler explanations of peptide data than positing the existence of very short, novel protein-coding genes.

Taking this a step further, we extended our previously-described method to allow it to evaluate downstream ORFs, and we then applied it to all upstream and downstream ORFs that were reported in [8] as novel proteins, novel isoforms, or peptideins (89 in total). We also incorporated long-read RNA-seq data, which were not used in [11]. Our method searches for alternative splice variants that explain the peptide evidence used for ncORFs, requiring that each transcript yields a translated protein at least 80% as long as the reference protein. For both short and long-read splicing evidence, we required the alignment evidence to span >70bp on both sides of the splice site.

Using all GENCODE (release 39, the same version used in [11] and [8]) mRNAs with complete ORFs, we found that 51 of 76 u/uoORFs and 11 of 13 d/doORFs had short and/or long read splicing evidence supporting the presence of protein-coding exons within these ncORF regions. In total, we found RNA-seq evidence for 229 exon candidates contained in 62 different ncORFs (both upstream and downstream combined). Thus for many ncORFs outside of the Tier 1A group from Deutsch *et al*., we found RNA-seq evidence supporting a much simpler explanation for the peptide evidence.

## Discussion

Many papers over the past 25 years have attempted to document the existence of new human genes, but with few exceptions, the human gene count has slowly declined as our understanding of the genome has improved. The number of human protein-coding genes currently hovers around 20,000, although major human gene catalogs (and annotation experts) have not yet converged on a precise estimate [2]. Two recent consortium-based efforts, including the one discussed here, have attempted to find new genes in a collection of 7,624 non-canonical open-reading frames [7, 8], most of which have experimental evidence that at least part of the ORF is being translated. The more recent Deutsch *et al*. study highlighted 137 “peptideins” as worthy of serious consideration for status as novel genes, and it ranked 19 of these candidates as particularly likely or even already confirmed as protein-coding genes.

The principal evidence for most of the claimed novel proteins is evidence that certain peptides appear to be translated from mRNA transcripts. We note that under the widely-accepted ribosome scanning model, the ribosome normally begins translating at the first ATG it encounters, but it is also well-established that some genes are translated at subsequent start codons [18]. Many of the top-ranking ncORFs in Deutsch *et al*. have start codons that occur upstream of the canonical, annotated start codon, providing a ready explanation for why the ribosome sometimes begins translation at those loci. Multiple studies have shown that translational noise from non-functional start codons is pervasive and often appears in both mass spectrometry and ribosome profiling data [17]. Thus one must be careful about inferring that a new gene has been discovered based primarily (or even partially) on evidence of translation.

The fact that only 0.25% of the original ncORFs made it into the highest tier of confidence illustrates how difficult it is to find new protein-coding genes that have gone undiscovered in the decades-long effort to catalog all human genes. Here we looked closely at each of the high-confidence candidates, and found serious problems with all 19 of them. Although three of the candidates are indeed genes, all three were previously known and published, and their inclusion in the collection of 7,264 ncORFs appears to have been an error. The other 16 candidates can credibly be explained without hypothesizing the existence of entirely new human proteins, for reasons that we detailed above. Notably, for the six candidates for which we identified possible alternative splice isoforms, the peptide evidence presented by Deutsch *et al*. could also be explained by translation of the alternative transcripts. Our findings suggest that the evidence presented by the TransCODE Consortium does not establish the existence of novel human protein-coding genes among these candidates, and that many can be explained by previously known genes, alternative splicing, or other forms of translation. There may indeed be a small number of novel human protein-coding genes yet to be discovered, but the large-scale analysis of ncORFs by the TransCODE Consortium has not, in our assessment, yielded any compelling examples.

## Data availability

The code for constructing transcripts based on RNA-seq evidence is publicly available on GitHub (https://github.com/haydenji0731/novex). The version used to generate all results in this paper is archived on Zenodo (https://doi.org/10.5281/zenodo.24352464). The GTF files containing the transcripts described in this paper, along with BED files of RNA-seq validated putative coding exons in peptidein regions, are archived on Zenodo (https://doi.org/10.5281/zenodo.24355222). The archive also includes the input files used for our analysis and a README.md file describing how to reproduce the results.

## Acknowledgements

This work was supported in part by NIH grants R01-HG006677 and R35-GM130151 to SLS.

---

**Supplementary Figure S1.**
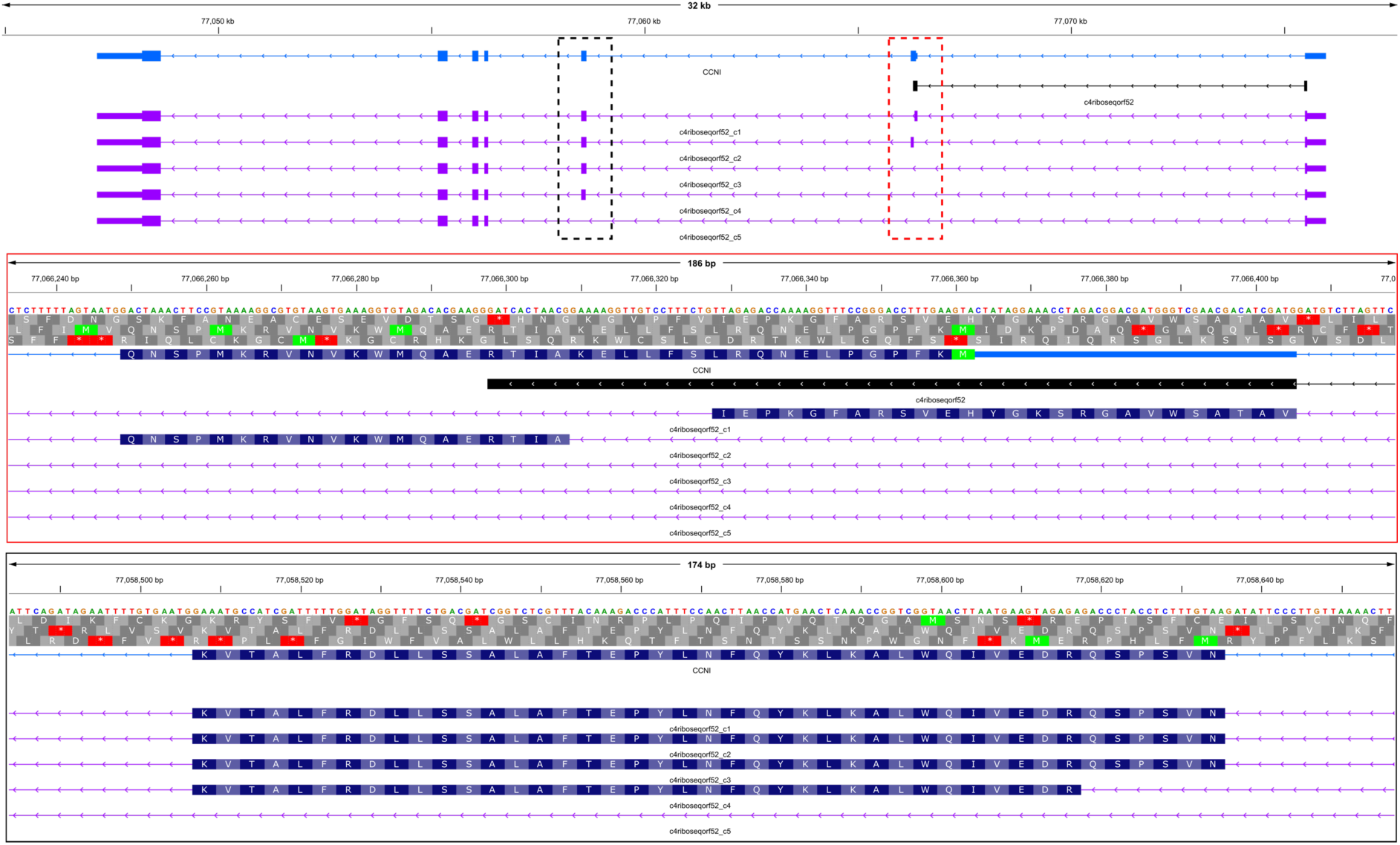
Structure of the CCNI gene, showing five alternative splicing variants that avoid the stop codon used by the ncORF c4riboseqorf52. The top panel shows the canonical (MANE) transcript in blue, the ncORF in black, and the alternative transcripts in mauve. Middle and bottom panels (in red and black solid lined boxes) are magnified views of exon 2 and exon 3 respectively. The middle panel shows that two of the alternative transcripts include portions of exon 2, while the others skip it entirely.

**Supplementary Figure S2.**
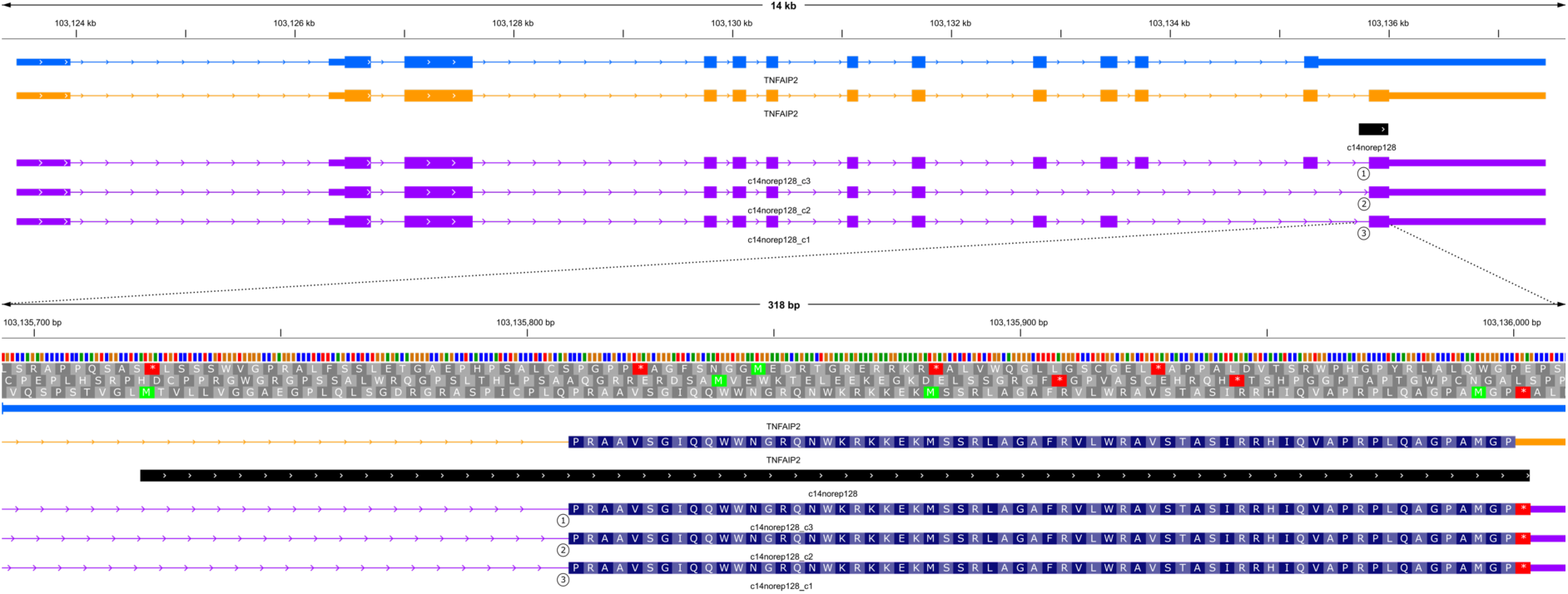
Structure of the TNFAIP2 gene, showing three distinct alternative splicing patterns that use a reference exon defined in the c14norep128 dORF region. The top panel shows the canonical (MANE) sequence in blue, a reference (RefSeq) isoform in yellow, the ncORF from Deutsch *et al*. in black, and the alternative transcripts described here in mauve. The bottom panel provides a closer view of the ncORF region, highlighting the coding exon that spans the final 64 amino acids of the open reading frame. This exon is shared by the RefSeq reference isoform (yellow) and three additional transcripts supported by RNA-seq evidence (mauve). Among the three RNA-seq-supported transcript constructs (labelled 1, 2, and 3), transcript 1 is supported by both short- and long-read RNA-seq data and is identical to the RefSeq reference isoform.

**Supplementary Table S1.**
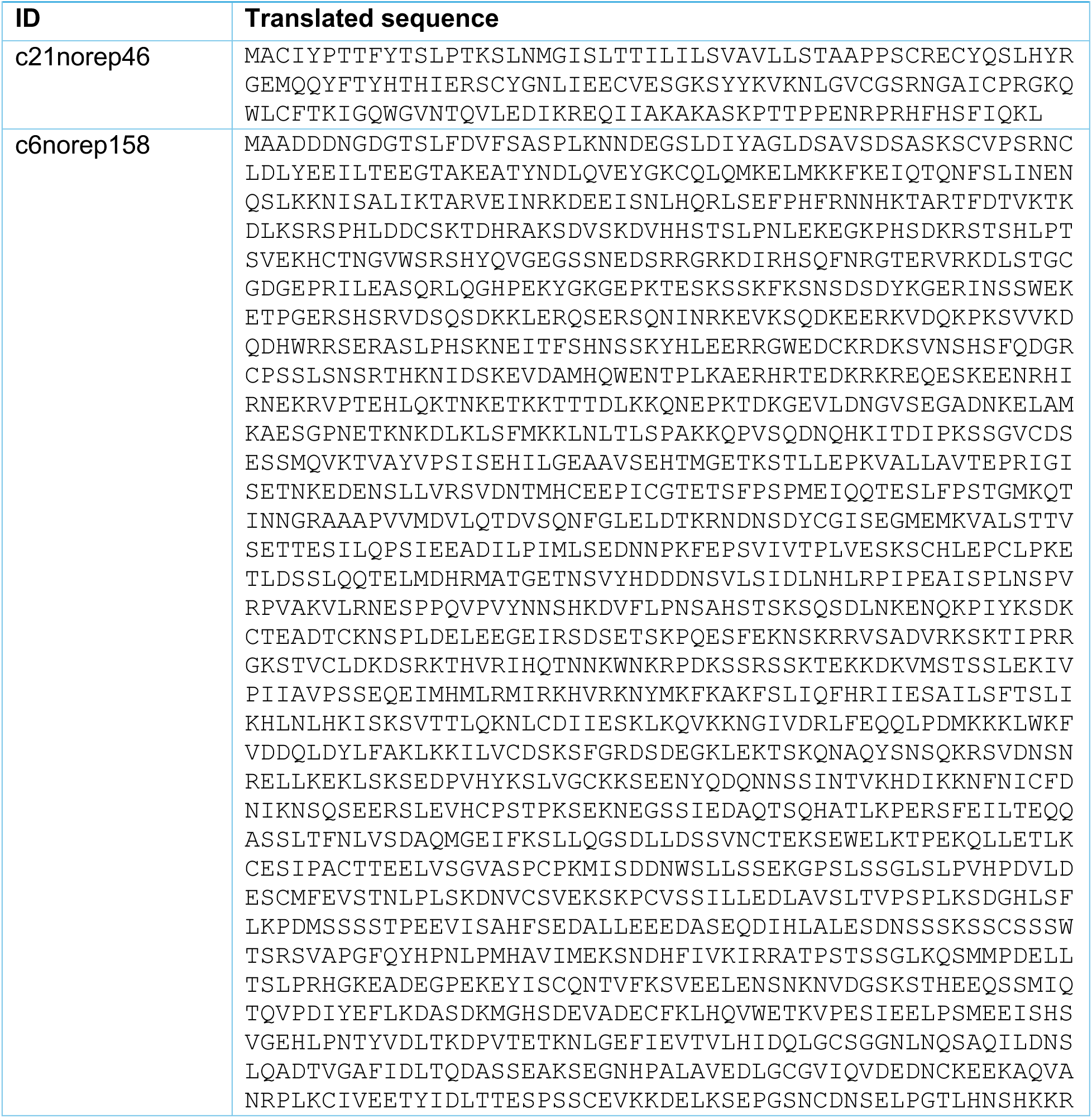
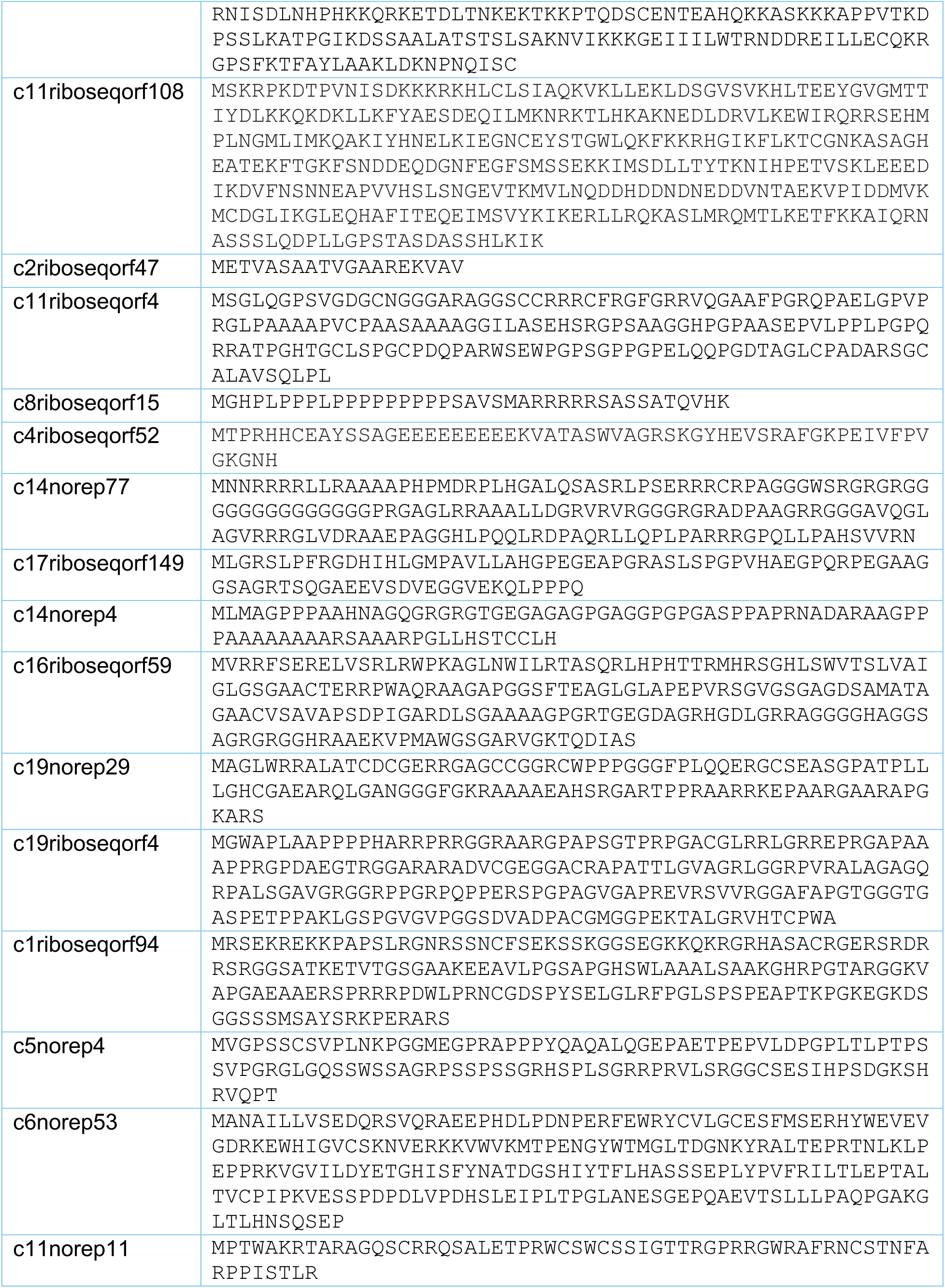
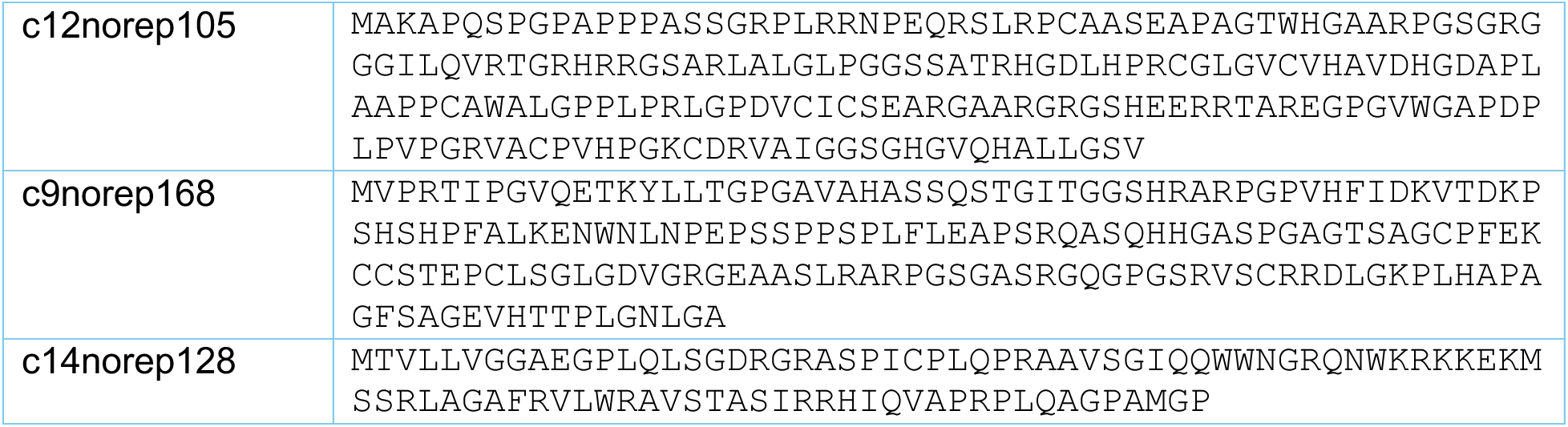
Amino acid sequences encoded by the ORFs shown in Table 1 and in Table S12 of Deutsch et al. [7].

| ID | Translated sequence |
|---|---|
| c21norep46 | MACIYPTTFYTSLPTKSLNMGISLTTILILSVAVLLSTAAPPSCRECYQSLHYRGEMQQYFTYHTHIERSCYGNLIEECVESGKSYYKVKNLGVCGSRNGAICPRGKQWLCFTKIGQWGVNTQVLEDIKREQIIAKAKASKPTTPPENRPRHFHSFIQKL |
| c6norep158 | MAADDDNGDGTSLFDVFSASPLKNNDEGSLDIYAGLDSAVSDSASKSCVPSRNCLDLYEEILTEEGTAKEATYNDLQVEYGKCQLQMKELMKKFKEIQTQNFSLINENQSLKKNISALIKTARVEINRKDEEISNLHQRLSEFPHFRNNHKTARTFDTVKTKDLKSRSPHLDDCSKTDHRAKSDVSKDVHHSTSLPNLEKEGKPHSDKRSTSHLPTSVEKHCTNGVWSRSHYQVGEGSSNEDSRRGRKDIRHSQFNRGTERVRKDLSTGCGDGEPRILEASQRLQGHPEKYGKGEPKTESKSSKFKSNSDSYKGERINSSWEKETPGERSHSRVDSQSDKKLERQSERSQNINRKEVKSQDKEERKVDQKPKSVVKDQDHWRRSERASLPHSKNEITFSHNSSKYHLEERRGWEDCKRDKSVNSHSFQDGRCPSSLSNSRTHKNIDSKEVDAMHQWENTPLKAERHRTEDKRKREQESKEENRHIRNEKRVPTEHLQKTNKETKKTTTDLKKQNEPKTDKGEVLDNGVSEGADNKELAMKAESGPNETKNKDLKLSFMKKLNLTLSPAKKQPVSQDNQHKITDIPKSSGVCDSESSMQVKTVAYVPSISEHILGEAAVSEHTMGETKSTLLEPKVALLAVTEPRIGISETNKEDENSLLVRSVDNTMHCEEPICGTETSFPSPMEIQQTESLFPSTGMKQTINNGRAAAPVVMDVLQTDVSQNFGLELDTKRNDNSDYCGISEGMEMKVALSTTVSETTESILQPSIEEADILPIMLSEDNNPKFEPSVIVTPLVESKSCHLEPCLPKETLDSSLQQTELMDHRMATGETNSVYHDDDNSVLSIDLNHLRPIPEAISPLNSPVRPVAKVLRNESPPQVPVYNNSHKDVFLPNSAHSTSKSQSDLNKENQKPIYKSDKCTEADTCKNSPLDELEEGEIRSDSETSKPQESFEKNSKRRVSADVRKSKTIPRRGKSTVCLDKDSRKTHVRIHQTNNKWNKRPDKSSRSSKTEKKDKVMSTSSLEKIVPIIAVPSSEQEIMHMLRMIRKHVRKNYMKFKAKFSLIQFHRIIESAILSFTSLIKHLNLHKISKSVTTLQKNLCDIIESKLKQVKKNGIVDRLFEQQLPDMKKKLWKFVDDQLDYLFAKLKKILVCDSKSFGRDSDEGKLEKTSKQNAQYSNSQKRSVDNSNRELLKEKLSKSEDPVHYKSLVGCKKSEENYQDQNNSSINTVKHDIKKNFNICFDNIKNSQSEERSLEVHCPSTPKSEKNEGSSIEDAQTSQHATLKPERSFEILTEQQASSLTFNLVSDAQMGEIFKSLLQGSDLLDSSVNCTEKSEWELKTPEKQLLETLKCESIPACTTEELVSGVASPCPKMISDDNWSLLSSEKGPSLSSGLSLPVHPDVLDESCMFEVSTNLPLSKDNVCSVEKSKPCVSSILLEDLAVSLTVPSPLKSDGHLSFLKPDMSSSSTPEEVISAHFSEDALLEEEDASEQDIHLALESDNSSSKSSCSSSWTSRSVAPGFQYHPNLPMHAVIMEKSNDHFIVKIRRATPSTSSGLKQSMMPDELLTSLPRHGKEADEGPEKEYISCQNTVFKSVEELENSNKNVDGSKSTHEEQSSMIQTQVPDIYEFLKDASDKMGHSDEVADECFKLHQVWETKVPESIEELPSMEEISHSVGEHLPNTYVDLTKDPVTETKNLGEFIEVTVLHIDQLGCSGGNLNQSAQILDNSLQADTVGAFIDLTQDASSEAKSEGNHPALAVEDLGCGVIQVDEDNCKEEKAQVANRPLKCIVEETYIDLTTESPSSCEVKKDELKSEPGSNCDNSELPGTLHNSHKKRRNISDLNHPHKKQRKETDLTNKEKTKKPTQDSCENTEAHQKKASKKKAPPVTKDPSSLKATPGIKDSSAALATSTSLSAKNVIKKKGEIIILWTRNDDREILLECQKRGPSFKTFAYLAAKLDKNPNQISC |
| c11riboseqorf108 | MSKRPKDTPVNISDKKKRKHLCLSIAQKVKLLEKLDSGVSVKHLTEEYGVGMTTIYDLKKQKDKLLKFYAESDEQILMKNRKTLHKAKNEDLDRVLKEWIRQRRSEHMPLNGMLIMKQAKIYHNELKIEGNCEYSTGWLQKFKKRHGIKFLKTCGNKASAGHEATEKFTGKFSNDDEQDGNFEGFSMSSEKKIMSDLLTYTKNIHPETVSKLEEEDIKDVFNSNNEAPVVHSLSNGEVTKMVLNQDDHDDNDNEDDVNTAEKVPIDDMVKMCDGLIKGLEQHAFITEQEIMSVYKIKERLLRQKASLMRQMTLKETFKKAIQRNASSSLQDPLLGPSTASDASSHLKIK |
| c2riboseqorf47 | METVASAATVGAAREKVAV |
| c11riboseqorf4 | MSGLQGPSVGDGCNGGGARAGGSCCRRRCFRGFGRRVQGAAFPGRQPAELGPVPRGLPAAAAPVCPAASAAAAGGILASEHSRGPSAAGGHPGPAASEPVLPPLPGPQRRATPGHTGCLSPGCPDQPARWSEWPGPSGPPGPELQQPGDTAGLCPADARSGCALAVSQLPL |
| c8riboseqorf15 | MGHPLPPPLPPPPPPPPPSAVSMARRRRRSASSATQVHK |
| c4riboseqorf52 | MTPRHHCEAYSSAGEEEEEEEEKVATASWVAGRSKGYHEVSRAFGKPEIVFPVGKGNH |
| c14norep77 | MNNRRRRLLRAAAAPHPMDRPLHGALQSASRLPSERRRCRPAGGGWSRGRGRGGGGGGGGGGGGPRGAGLRRAAALLDGRVRVRGGGRGRADPAAGRRGGGAVQGLAGVRRRGLVDRAAEPAGGHLPQQLRDPAQRLLQPLPARRRGPQLLPAHSVVRN |
| c17riboseqorf149 | MLGRSLPFRGDHIHLGMPAVLLAHGPEGEAPGRASLSPGPVHAEGPQRPEGAAGGSAGRTSQGAEEVSDVEGGVEKQLPPPQ |
| c14norep4 | MLMAGPPPAAHNAGQGRGRGTGEGAGAGPGAGGPGPGASPPAPRNADARAAGPPPAAAAAAAARSAAARPGLLHSTCCLH |
| c16riboseqorf59 | MVRRFSERELVSRLRWPKAGLNWILRTASQRLHPHTTRMHRSGHLSWVTSLVAIGLGSGAACTERRPWAQRAAGAPGGSFTEAGLGLAPEPVRSGVGSGAGDSAMATAGAACVSAVAPSDPIGARDLSGAAAAGPGRTGEGDAGRHGDLGRRAGGGGHAGGSAGRGRGGHRAAEKVPMAWGSGARVGKTQDIAS |
| c19norep29 | MAGLWRRALATCDCGERRGAGCCGGRCWPPPGGGFPLQQERGCSEASGPATPLLLGHCGAEARQLGANGGGFGKRAAAAEAHSRGARTPPRAARRKEPAARGAARAPGKARS |
| c19riboseqorf4 | MGWAPLAAPPPPHARRPRRGGRAARGPAPSGTPRPGACGLRRLGRREPRGAPAAAPPRGPDAEGTRGGARARADVCGEGGACRAPATTLGVAGRLGGRPVRALAGAGQRPALSGAVGRGGRPPGRPQPPERSPGPAGVGAPREVRSVVRGGAFAPGTGGGTGASPETPPAKLGSPGVGVPGGSDVADPACGMGGPEKTALGRVHTCPWA |
| c1riboseqorf94 | MRSEKREKKPAPSLRGNRSSNCFSEKSSKGGSEGKKQKRGRHASACRGERSRDRRSRGGSATKETVTGSGAAKEEAVLPGSAPGHSWLAAALSAAKGHRPGTARGGKVAPGAEAAERSPRRRPDWLPRNCGDSPYSELGLRFPGLSPSPEAPTKPGKEGKDSGGSSSMSAYSRKPERARS |
| c5norep4 | MVGPSSCSVPLNKPGGMEGPRAPPPYQAQALQGEPAETPEPVLDPGPLTLPTPSSVPGRGLGQSSWSSAGRPSSPSSGRHSPLSGRRPRVLSRGGCSESIHPSDGKSHRVQPT |
| c6norep53 | MANAILLVSEDQRSVQRAEEPHDLPDNPERFEWRYCVLGCESFMSERHYWEVEVGDRKEWHIGVCSKNVERKKVWVKMTPENGYWTMGLTDGNKYRALTEPRTNLKLPEPPRKVGVILDYETGHISFYNATDGSHIYTFLHASSSEPLYPVFRILTLEPTALTVCPIPKVESSPDPDLVPDHSLEIPLTPGLANESGEPQAEVTSLLLPAQPGAKGLTLHNSQSEP |
| c11norep11 | MPTWAKRTARAGQSCRRQSALETPRWCSWCSSIGTTRGPRRGWRAFRNCSTNFARPPISTLR |
| c12norep105 | MAKAPQSPGPAPPPASSGRPLRRNPEQRSLRPCAASEAPAGTWHGAARPGSGRGGGILQVRTGRHRRGSARLALGLPGGSSATRHGDLHPRCGLGVCVHAVDHGDAPLAAPPCAWALGPPLPRLGPDVCICSEARGAARGRGSHEERRTAREGPGVWGAPDPLPVPGRVACPVHPGKCDRVAIGGSGHGVQHALLGSV |
| c9norep168 | MVPRTIPGVQETKYLLTGPGAVAHASSQSTGITGGSHRARPGPVHFIDKVTDKPSHSHPFALKENWNLNPEPSSPPSPLFLEAPSRQASQHHGASPGAGTSAGCPFEKCCSTEPCLSGLGDVGRGEAASLRARPGSGASRGQGPGSRVSCRRDLGKPLHAPAGFSAGEVHTTPLGNLGA |
| c14norep128 | MTVLLVGGAEGPLQLSGDRGRASPICPLQPRAAVSGIQQWWNGRQNWKRKKEKMSSRLAGAFRVLWRAVSTASIRRHIQVAPRPLQAGPAMGP |

